# CoExp Web, a web tool for the exploitation of co-expression networks

**DOI:** 10.1101/2020.06.29.176057

**Authors:** Sonia García-Ruiz, Ana L. Gil-Martínez, Alejandro Cisterna, Federico Jurado-Ruiz, Regina R. Reynolds, NABEC (North America Brain Expression Consortium), Mark R. Cookson, John Hardy, Mina Ryten, Juan A. Botía

## Abstract

**Summary:** This paper presents the CoExp Web tool. CoExp is a software tool and a Web resource that fosters research on co-expression network models by providing a highly usable means to exploit and share multiple network models.

**Availability and Implementation:** the Web resource is accessible at https://snca.atica.um.es/coexp/. The software at the front and back-ends is accessible at https://github.com/SoniaRuiz/CoExp_Web.

**Contact:** (Sonia García-Ruíz)

## Introduction

Gene co-expression network analysis has been widely used to identify biologically important patterns in gene expression in a hypothesis-free and genome-wide manner^1–4^. The driving principle behind co-expression network analysis is that genes whose expression levels are highly correlated, are also likely to share functional and biological relationships^5,6^. Thus, co-expression networks (CEN) are models of how genes cluster together into modules of highly co-expressed genes, generally by using graph-based approaches to reveal their similarities^7^. A CEN is useful as a way to describe gene-gene relationships in a given gene expression profile which represent a given condition (e.g. a tissue, a specific cell type or a group of cases and controls). But CENs can also be useful when used for annotation of external gene sets. For example, (1) by looking at whether the genes cluster together in the network, (2) by identifying new genes co-expressed with the input genes, or (3) specific cell type markers clustering together with those genes.

Normally, managing CEN models is work intensive, and becomes highly complex when multiple CENs are being explored. In these cases, a Web based application integrating all those CENs, becomes the optimum solution. We present here CoExp, accessible at https://snca.atica.um.es/coexp/. CoExp is a Web application to increase the usability and accessibility of CENs. CoExp offers 109 different CENs created with the CoExpNets R package^7^, based on the widely used WGCNA R package^8^ (see a, Figure 1).

**Figure 1:**
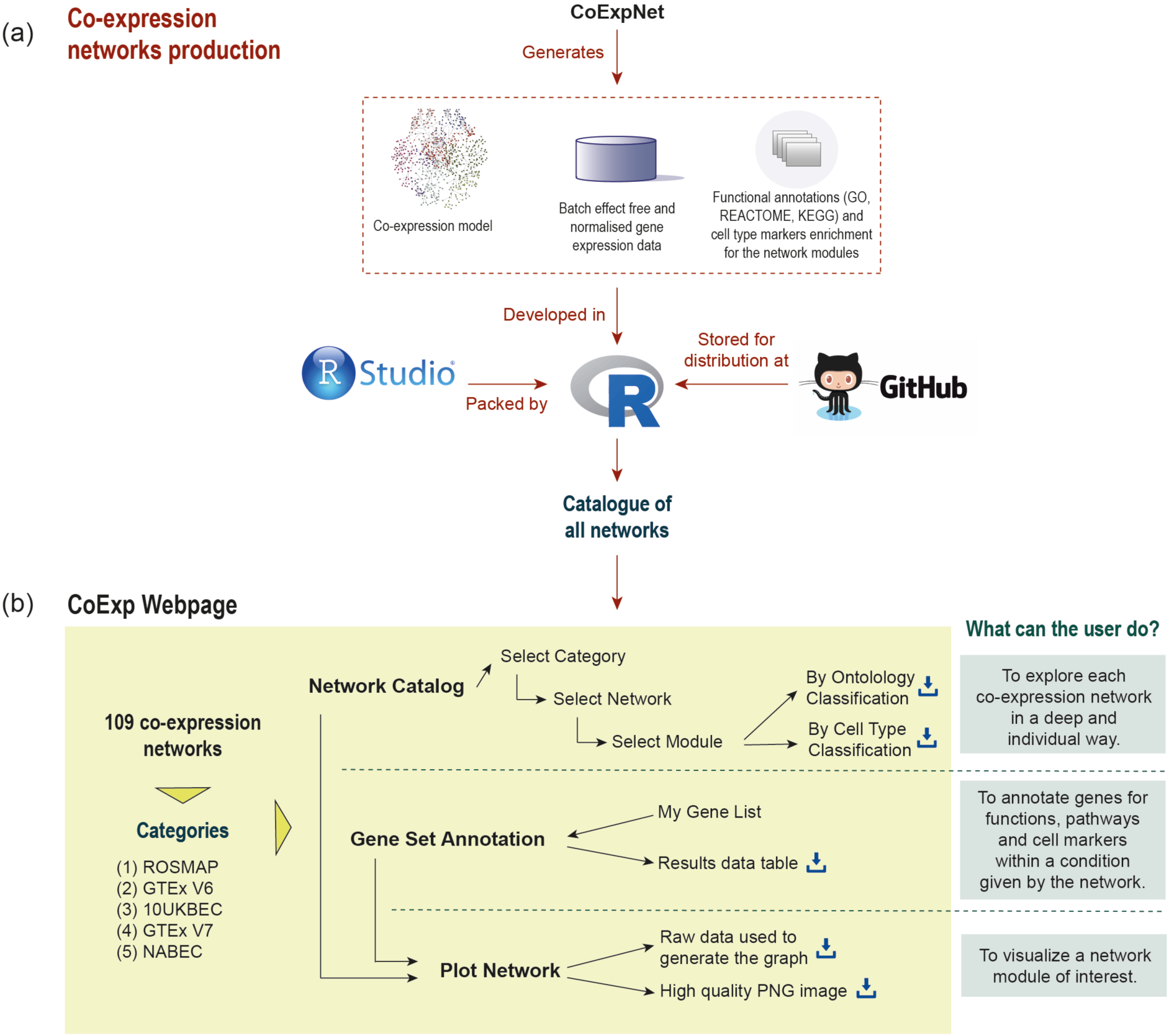
(a) All 109 networks in CoExp were generated with CoExpNets R package for co-expression network generation and annotation. Networks and annotations are organised into R packages (i.e. one package for each network category) and stored in GitHub. (b) There are five network categories currently available at CoExp. All of them can be browsed, used for gene set annotation and plotted by gene modules.

The network catalogue is organised into four different network groups: (1) the Religious Orders Study and Memory and Aging Project (ROSMAP) ^9–11^ composed of four CENs derived from RNAseq data on post-mortem human frontal cortex originating from control individuals, as well as those with cognitive impairment and Alzheimer’s disease; (2) The Genotype-Tissue Expression project (GTEx, V6 and V7^12^) composed of two suites of CENs derived from RNAseq data on 47 post-mortem control human tissue samples; (3) the UK Brain Expression Consortium project (UKBEC) ^2,13^ composed of ten microarray-based CENs derived from post-mortem control human brain tissue; (4) the North America Brain Expression Consortium (NABEC)^14^, composed of one CEN derived from RNAseq data on post-mortem control human frontal cortex (see b, Figure 1).

### Network Catalogue Browser

The user can become familiar with the CEN models available by browsing, and downloading, the catalogue through the ‘Network Catalogue’ tab. The user selects a network category, and then a specific network. Finally, the user decides between two different views of the network: the ‘Ontology Classification’ or the ‘Cell Type Classification’. The ‘Ontology Classification’ view displays an active data table in which each module (i.e. gene cluster) from the selected network occupies one row. The columns provide summarized information about annotation terms enriched for the genes in the modules. Those terms come from an enrichment analysis performed with gProfileR ^15^, which incorporates data from well recognised ontologies, including Gene Ontology^16^, REACTOME^17^ and KEGG^18^. The ‘Cell Type Classification’ view, displays an active data table in which the rows correspond now to sets of gene markers of brain cell types tested for enrichment (Fisher’s Exact test) and each module occupies a column. Each cell within the table contains the Bonferroni corrected p-value for the enrichment of the corresponding cell markers within the module. In both view types, the raw data behind the tables is easily downloadable.

### Gene Set Annotation

CENs are often used to annotate a gene set of interest, in the context of a specific condition (e.g. a tissue of interest). For those users, CoExp offers the ‘Gene Set Annotation’ tab. Thanks to this function the user can investigate whether the gene set is enriched within a single or multiple CEN modules, across all the co-expression networks from amongst those available in the catalogue. In this way, those genes can be annotated based on how they are distributed across the network modules and their biological context explored within just a couple of clicks. The user is supplied a results table in which each row relates to a gene of interest which has been successfully found in any of the modules belonging to the network or networks selected. The columns provide information on the module in which the gene has been found, including the statistical significance of the overlap between the input genes and the genes int the module and a brief description of the module’s function based on the its top five GO terms. This analysis outcome can easily be downloaded as an excel file.

### Plot Network

Once the user has decided which network is of interest, and which module within the network requires detailed visualization, the corresponding gene graph can be plotted. The third tab, ‘Plot Network’, enables the graph-based visualisation of the genes within a network module of interest, whether identified by browsing through the catalogue or because the user’s gene set of interest significantly clusters within a module. The ‘Plot Network’ tab generates an interactive directed graph formed by the hub (i.e. most important) genes within a module. The user can select how many of the most relevant genes will appear within the plot. The resulting plot is interactive in the sense that it can be zoomed, rotated and the direct neighbours of any gene highlighted by just clicking on the gene of interest. Both the raw data displayed in the graph and a high quality PNG image are available for download.

## Conclusions

CoExp Webpage is a web platform for the exploitation of co-expression networks. CoExp currently offers 109 CENs focused on brain transcriptomics with plans to expand its scope. It is a powerful, easy to use and innovative tool for gene set annotation across a variety of brain-specific transcriptomic datasets. CoExp makes CENs easily shareable with the scientific community. Everything is downloadable in CoExp, including the CENs themselves and the front-and back-end software so any research laboratory can construct their own CoExp web site. This makes it a powerful tool for the wider research community interested in producing, using or sharing co-expression models to support their research.

## Data availability

All co-expression models available through CoExp and their respective annotations can be downloaded from the CoExp application, accessible from https://snca.atica.um.es/coexp/. Alternatively, gene expression profiles and networks created from those expression profiles, for all networks available at CoExp can be obtained at GitHub in the form of R packages, one package per co-expression network family. All the packages and networks were created using CoExpNets software, based on WGCNA, accessible at https://github.com/juanbot/CoExpNets. GTEx networks and expression profiles can be downloaded from https://github.com/juanbot/CoExpGTEx. ROSMAP networks and expression profiles can be downloaded from https://github.com/juanbot/CoExpROSMAP. UKBEC networks and expression profiles can be downloaded from https://github.com/juanbot/CoExp10UKBEC. Finally, NABEC network and expression profiles can be obtained through https://github.com/juanbot/CoExpNABEC. The back and front ends code of the CoExp Web application is fully available for download on GitHub at https://github.com/SoniaRuiz/CoExp_Web.

## Acknowledgements

Study data to create the ROSMAP networks were provided by the Rush Alzheimer’s Disease Center, Rush University Medical Center, Chicago. Data collection was supported through funding by NIA grants P30AG10161, R01AG15819, R01AG17917, R01AG30146, R01AG36836, U01AG32984, U01AG46152, the Illinois Department of Public Health and the Translational Genomics Research Institute.

## References

1. de la Torre-Ubieta, L. et al. The Dynamic Landscape of Open Chromatin during Human Cortical Neurogenesis. Cell 172, 289-304.e18 (2018).

2. Forabosco, P. et al. Insights into TREM2 biology by network analysis of human brain gene expression data. Neurobiol. Aging 34, 2699–2714 (2013).

3. Bettencourt, C. et al. White matter DNA methylation profiling reveals deregulation of HIP1, LMAN2, MOBP, and other loci in multiple system atrophy. Acta Neuropathol. (Berl.) (2019) doi: 10.1007/s00401-019-02074-0.

4. UK Brain Expression Consortium (UKBEC) et al. Frontotemporal dementia: insights into the biological underpinnings of disease through gene co-expression network analysis. Mol. Neurodegener. 11, (2016).

5. Xuelian Ma. Co-expression Gene Network Analysis and Functional Module Identification in Bamboo Growth and Development. frontiers in Genetics (2018) doi: https://doi.org/10.3389/fgene.2018.00574.

6. Mohammad Reza Bakhtiarizadeh. Weighted Gene Co-expression Network Analysis of Endometriosis and Identification of Functional Modules Associated With Its Main Hallmarks. frontiers in Genetics (2018) doi: https://doi.org/10.3389/fgene.2018.00453.

7. An additional k-means clustering step improves the biological features of WGCNA gene co-expression networks. BMC Systems Biology (2017) doi: https://doi.org/10.1186/s12918-017-0420-6.

8. Langfelder, P. & Horvath, S. WGCNA: an R package for weighted correlation network analysis. BMC Bioinformatics 9, 559 (2008).

9. A. Bennett, D., A. Schneider, J., Arvanitakis, Z. & S. Wilson, R. Overview and Findings from the Religious Orders Study. Curr. Alzheimer Res. 9, 628–645 (2012).

10. A. Bennett, D. et al. Overview and Findings from the Rush Memory and Aging Project. Curr. Alzheimer Res. 9, 646–663 (2012).

11. De Jager, P. L. et al. A multi-omic atlas of the human frontal cortex for aging and Alzheimer’s disease research. Sci. Data 5, 180142 (2018).

12. The GTEx Consortium et al. The Genotype-Tissue Expression (GTEx) pilot analysis: Multitissue gene regulation in humans. Science 348, 648–660 (2015).

13. UK Brain Expression Consortium et al. Genetic variability in the regulation of gene expression in ten regions of the human brain. Nat. Neurosci. 17, 1418–1428 (2014).

14. Dillman, A. A. et al. Transcriptomic profiling of the human brain reveals that altered synaptic gene expression is associated with chronological aging. Sci. Rep. 7, (2017).

15. Reimand, J., Kull, M., Peterson, H., Hansen, J. & Vilo, J. g:Profiler—a web-based toolset for functional profiling of gene lists from large-scale experiments. Nucleic Acids Res. 35, W193–W200 (2007).

16. The Gene Ontology Consortium. Expansion of the Gene Ontology knowledgebase and resources. Nucleic Acids Res. 45, D331–D338 (2017).

17. Fabregat, A. et al. The Reactome Pathway Knowledgebase. Nucleic Acids Res. 46, D649–D655 (2018).

18. Kanehisa, M., Sato, Y., Kawashima, M., Furumichi, M. & Tanabe, M. KEGG as a reference resource for gene and protein annotation. Nucleic Acids Res. 44, D457–D462 (2016).

